# Intranuclear niche: actin-tail mediated nuclear entry by intracellular pathogens

**DOI:** 10.64898/2026.08.23.746248

**Authors:** Hoi Ching Cheung, Patricia Reist Iscar, Miro Thorsten Wilhelm Plum, Marek Basler

## Abstract

Intracellular pathogens localize to various niches in the host cells to avoid immune detection. However, very little is known about bacteria that enter the host nuclei. Here we report that *Burkholderia thailandensis*, a facultative intracellular pathogen, can enter eukaryotic nuclei and replicate. Nuclear invasion events were rare, occurring 1 in 500-1,000 infected cells, and inhibition of cell division further reduced the frequency of these events. Moreover, we show that nuclear entry requires actin tail motility, although it is independent of other virulence factors such as the Type III Secretion System, Type VI Secretion System-5, and flagella motility. Inactivation of actin tail motility by deleting *bimA* or inhibition of actin polymerisation by cytochalasin D abolished nuclear entry. Surprisingly, we observed that accumulation of *B. thailandensis* in the nucleus activated assembly of the Type VI Secretion Systems-5. We further show that *Shigella flexneri* also enters nucleus in an actin polymerization dependent mechanism. Together, we show that actin tail forming intracellular pathogens occasionally localize to the nucleus, and while this largely requires host cell division, it may provide pathogens with a protective niche in certain mitotically active cells, such as skin, gut or epithelial cells.

## Introduction

To avoid host innate immune detection, intracellular pathogens often occupy specific niches and modify the host environment. Most intracellular pathogens replicate either in the cytoplasm or in vacuoles. For example, *Salmonella* mostly resides in Salmonella-containing vacuoles (SCV), and *Listeria* and *Shigella* spp. replicate in the host cytosol(1–3). In rare cases, some bacteria such as *Holospora* spp., *Candidatus* spp., *Rickettsia rickettsii* and *Rickettsia bellii* have been reported to localize to the intranuclear compartment(4–9). Although *Holospora* spp. and *Candidatus Nucleicultrix amoebiphila* are described as symbionts in the nucleus of amoeba, the intranuclear niche is not well characterized in the context of mammalian infections(10,11). A facultative intracellular pathogen - *Burkholderia pseudomallei* has been reported to localize to the nucleus of infected human lung and guinea pig spleen tissues. It is speculated that intranuclear bacteria contributes to latent infections and disease relapse(12).

*Burkholderia* species are gram-negative, rod-shaped bacteria present in a variety of environments, mostly in soil and water, but some are opportunistic pathogens(13,14). *B. thailandensis*, *B. pseudomallei* and *B. mallei* are classified as virulent species and are genetically highly similar with over 85% of the genome conserved(15). *B. pseudomallei* causes melioidosis in humans, which is a disease with mortality rate as high as 40% even with the appropriate antibiotic treatment(16,17). A modelling study estimated that there are around 165,000 cases of melioidosis in humans per year worldwide(17,18). In particular, *B. pseudomallei* is able to cause latent and relapsing infections in the host, and thus it is classified as a biosafety level 3 and Tier 1 select agent(19,20). *B. thailandensis* is often used as a surrogate biosafety level 2 organism(21), yet there are also reports of *B. thailandensis* causing severe lung and wound infections in humans(22–24).

Upon entry into host cell in a membrane-bound vacuole, the *Burkholderia* utilise the Type III Secretion System (T3SS) to escape from the endosome before phagolysosomal fusion(25). The bacterium replicates in the host cytosol and either swims with an intracellular flagella, or hijacks host actin to form actin tails(26–30). Together with their Type VI Secretion System 5 (T6SS-5), the bacterium can spread to neighbouring cells by stimulating the formation of multinucleated giant cells (MNGC) or by lysing protrusions for direct cell-to-cell spread (20,31–33).

One of the key features of *B. pseudomallei* infections is its ability to remain in the host for a prolonged period of time(34,35). It is unclear how *Burkholderia* species cause latent infections, and no virulence factors were shown to play a role in latency. However, virulence factors such as the capsular polysaccharide and the formation of MNGCs have been suggested(36,37). MNGCs have been found inside of granuloma-like lesions formed during *Burkholderia* infections(38,39).

In this study, we found that *B. thailandensis* localize to host cell nuclei. While this is rare, up to 1 in 500-1000 infected cells, the bacteria are able to replicate within the nucleus and activated assembly of its T6SS-5 when accumulated to a high density. *B. thailandensis* invades host nuclei using actin-based motility predominantly during host cell division. Interestingly, *S. flexneri*, another intracellular pathogen capable of actin-based motility, also invades the nucleus. Overall, our results reveal actin-based entry to the intranuclear niche, which is potentially significant for survival and spread within the host.

## Results

### *Burkholderia thailandensis* are present and can replicate in eukaryotic cell nuclei

When observing the intracellular life cycle of *B. thailandensis* in A549 lung cells, we noticed an accumulation of bacteria in eukaryotic cell nuclei visualized with Hoechst 33342 (Figure 1A, S1, Video S1). To ensure *B. thailandensis* were inside rather than on the surface of the nucleus, we collected a 15.4 μm z-stack image of *B. thailandensis* co-localizing with host cell nuclei at 12:30 hpi (Figure 1B). At early stage of nuclear invasion (11:30 hpi), only a few bacteria were found within the nuclei. The bacteria were non-motile, replicated (15:30 hpi), and filled the host cell nucleus at later stages (18:30 hpi).

**Figure 1.**
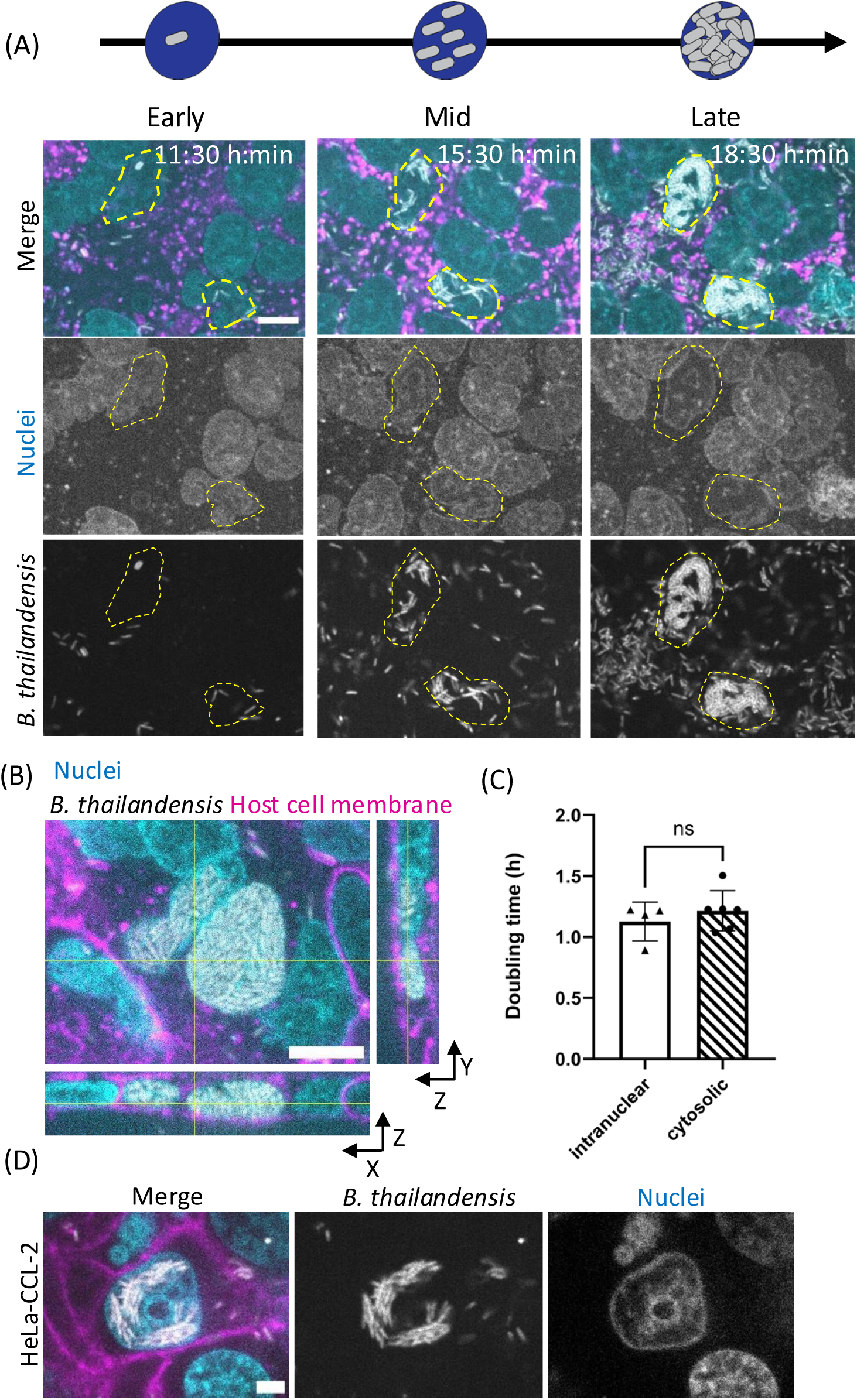
*Burkholderia thailandensis* invades eukaryotic cell nuclei. **(A)** Time-lapse imaging of A549 lung cells infected with *B. thailandensis* expressing TssB-5-mScarlet-I (grey) at MOI of 50 in the early, mid and late stages of infection. Infection was performed in Opti-MEM^TM^ supplemented with 300 μg/mL of kanamycin. Host cell membrane was stained with CellMask DeepRed (magenta) and nuclei stained with Hoechst 33342 (cyan). Nuclei containing *B. thailandensis* are outlined with yellow dotted lines. Scale bar represents 10 µm. **(B)** Single z-slice image of a 15.4 μm z-stack image of *B. thailandensis* co-localizing with host cell nuclei. A549 lung cells were infected with *B. thailandensis* expressing TssB-5-mScarlet-I (grey) at MOI of 50. Images were obtained at 12:30 hpi. Host cell membrane was stained with CellMask DeepRed (magenta) and nuclei stained with Hoechst 33342 (cyan). Orthogonal views covering the whole 15.4 μm z-stack are displayed. Scale bar represents 10 µm. **(C)** Doubling time of intranuclear bacteria and cytosolic bacteria calculated by approximating a Malthusian exponential growth curve. An unpaired *t*-test was performed (ns = not significant). **(D)** Nuclear invasion event in HeLa CCL-2 cells. HeLa CCL-2 cells were infected with *B. thailandensis* expressing TssB-5-mScarlet-I (grey) at MOI of 100 and images were taken at 14 hpi. Host cell membrane was stained with CellMask DeepRed (magenta) and nuclei stained with Hoechst 33342 (cyan). Scale bar represents 5 µm.

To compare the growth of intranuclear and cytosolic *B. thailandensis*, we quantified their doubling time (Figure 1C). To simplify tracking of cytoplasmic bacteria, an actin-tail motility mutant *B. thailandensis* Δ*bimA* was used. A monolayer of A549 cells were infected and confocal images were taken at 10-minute intervals. Bacterial replication was followed for bacteria in 4 infected nuclei and the cytosol of 6 infected cells for at least 10 hours. The doubling time was calculated by approximating a Malthusian exponential growth curve. The doubling time were on average 1.13 hours and 1.21 hours for intranuclear (*bimA*-positive) and cytosolic (*bimA*-negative) bacteria respectively. This indicates that the host nucleus supports unrestricted growth of *B. thailandensis*.

To exclude that this observation is specific for A549 cell line we also infected a monolayer of HeLa-CCL-2. In 4,471 infected HeLa cells over two biological replicates, we observed 8 nuclear invasion events, which is comparable to what is observed in A549 cells (Figure 1D). These results indicate that nuclear invasion occurs across multiple cell types.

### *B. thailandensis* accumulated in confined compartments assemble the T6SS-5

Recently, T6SS-5 was shown to preferably assemble in protrusions(33), we wondered if T6SS-5 also assembles when the bacterium is inside of the nucleus. To investigate this, *B. thailandensis* expressing the sheath protein TssB-5 tagged with mScarlet-I were imaged at 30 second intervals for 15 minutes to visualize T6SS-5 assemblies identified as dynamically appearing foci of TssB5-mScarlet-I (Figure 2A, Video S2). We calculated the percentage of nucleus area occupied with bacteria throughout the imaging time period and discovered that T6SS-5 assemblies were only observed when the bacteria accumulated at late time-points (Figure 2B). At 7:30 hpi, when only ∼10% of the nucleus was occupied with bacteria, no T6SS-5 assemblies were observed. However, at 11:30 hpi, when the bacteria had accumulated in over ∼60% of the nucleus, T6SS-5 assemblies could be detected. Within the same infected cell, we detected no T6SS-5 assemblies in the cytosolic bacteria at 11:30 hpi (Figure 2C). In addition, when there were two nuclear invasion events with different bacterial densities in the same MNGC, we noticed that only the intranuclear bacteria that accumulated to high density assembled T6SS-5 (Figure 2D, Video S2).

**Figure 2.**
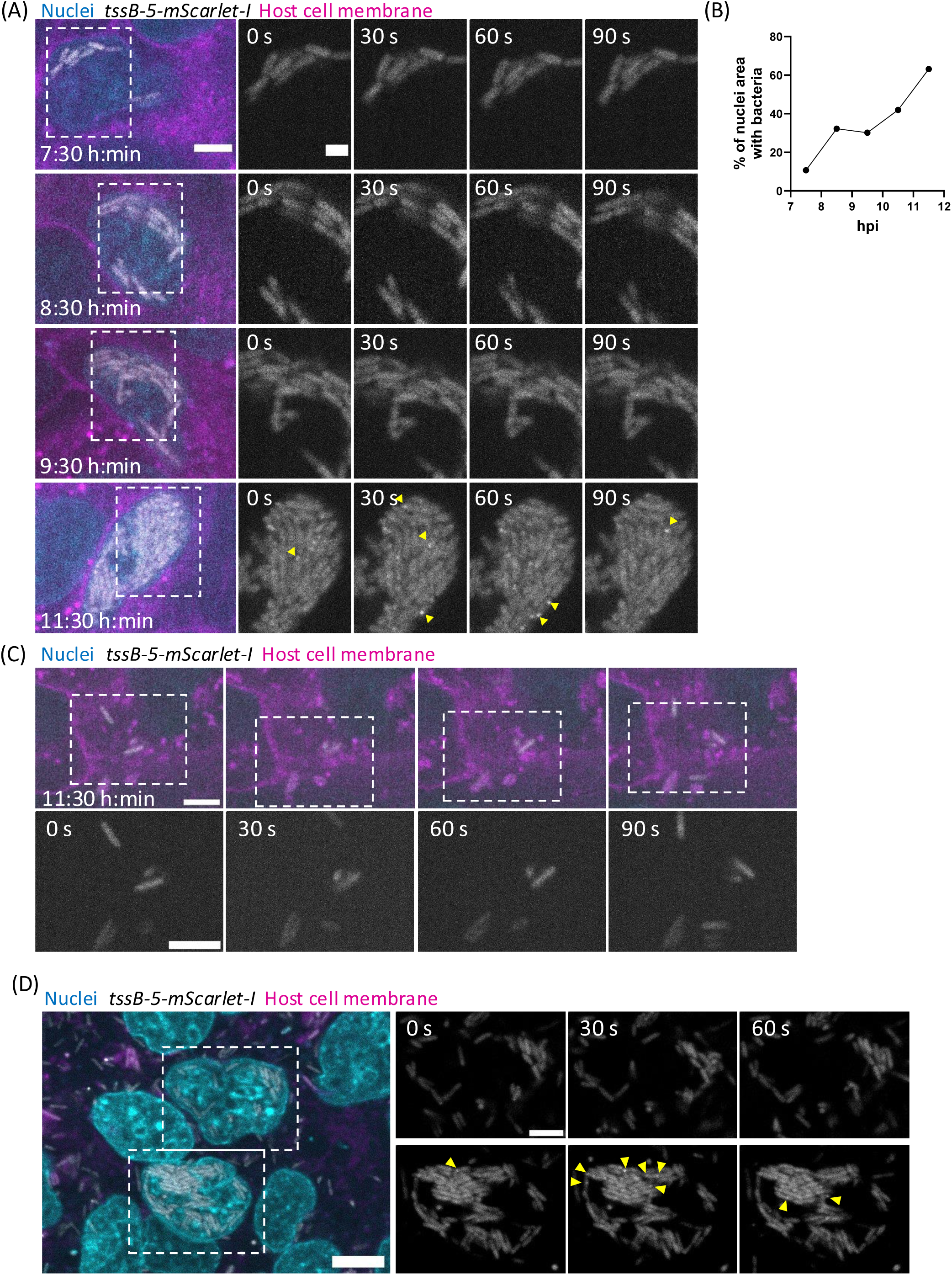
T6SS-5 assembles in *B. thailandensis* accumulated in the nucleus or cytoplasm. **(A)** T6SS-5 assemblies observed for intranuclear *B. thailandensis.* A549 cells were infected with *B. thailandensis* expressing TssB-5-mScarlet-I (grey) at MOI of 50. Infection was performed in Opti-MEM^TM^ supplemented with 300 μg/mL of kanamycin. Confocal images were taken at 30 second intervals for a duration of 15 minutes at every hour post-infection. Yellow arrows indicate T6SS-5 assemblies. Scale bar on the larger image represents 5 μm and 2 μm in the magnified images. **(B)** Percentage of host nuclei area covered with bacteria over time. Time-lapse images from (A) was analysed with ImageJ. Bacterial fluorescence intensity were thresholded and particle size were analysed to obtain the area occupied by bacteria. Host nuclei were manually segmented and area was measured. **(C)** Cytosolic *B. thailandensis* from the same infected cell as (A) at 11:30 hpi. Scale bars represent 5 μm. **(D)** T6SS-5 assemblies observed for intranuclear *B. thailandensis.* A549 cells were infected with *B. thailandensis* expressing TssB-5-mScarlet-I (grey) at MOI of 50. Confocal images were taken at 30 second intervals at 15:30 hpi. Yellow arrows indicate T6SS-5 assemblies. Scale bar on the larger image represents 10 μm and 5 μm in the magnified images.

To further explore if *B. thailandensis* assembles T6SS-5 when confined in compartments, we infected A549 cells with *B. thailandensis* Δ*bimA*, which was shown to accumulate in host cytosol(28,40–42) (Figure S2A). Indeed, when we imaged the Δ*bimA* mutant at 6:30 hpi at 30 seconds intervals, we observed no T6SS-5 assemblies. However, at 13 hpi, when the bacteria have accumulated inside the host cell cytosol, we observed that up to 70 % of the bacteria assembled T6SS-5 (Figure S2B, Video S3). This suggested that high bacterial density, regardless of compartment, promotes T6SS-5 assembly.

### Infected cells stop synthesizing new DNA

After confirming that the bacteria were able to replicate and assemble T6SS-5, we next examined whether cells with infected nuclei remained metabolically active by measuring EdU incorporation, a marker of ongoing DNA synthesis. EdU is a fluorescent thymidine analogue that is incorporated into newly synthesized DNA during S phase(43). A549 cell monolayers were infected and subjected to EdU incorporation analysis at 15 h post-infection (hpi), alongside uninfected control cells. All nuclei were identified with Hoechst 33342 staining, and we noted that only a subset of A549 cells incorporated EdU in the uninfected control (Figure 3A). Interestingly, we observed that infected cells generally do not synthesize new DNA, and that none of the 63 infected nuclei synthesized new DNA (Figure 3B).

**Figure 3.**
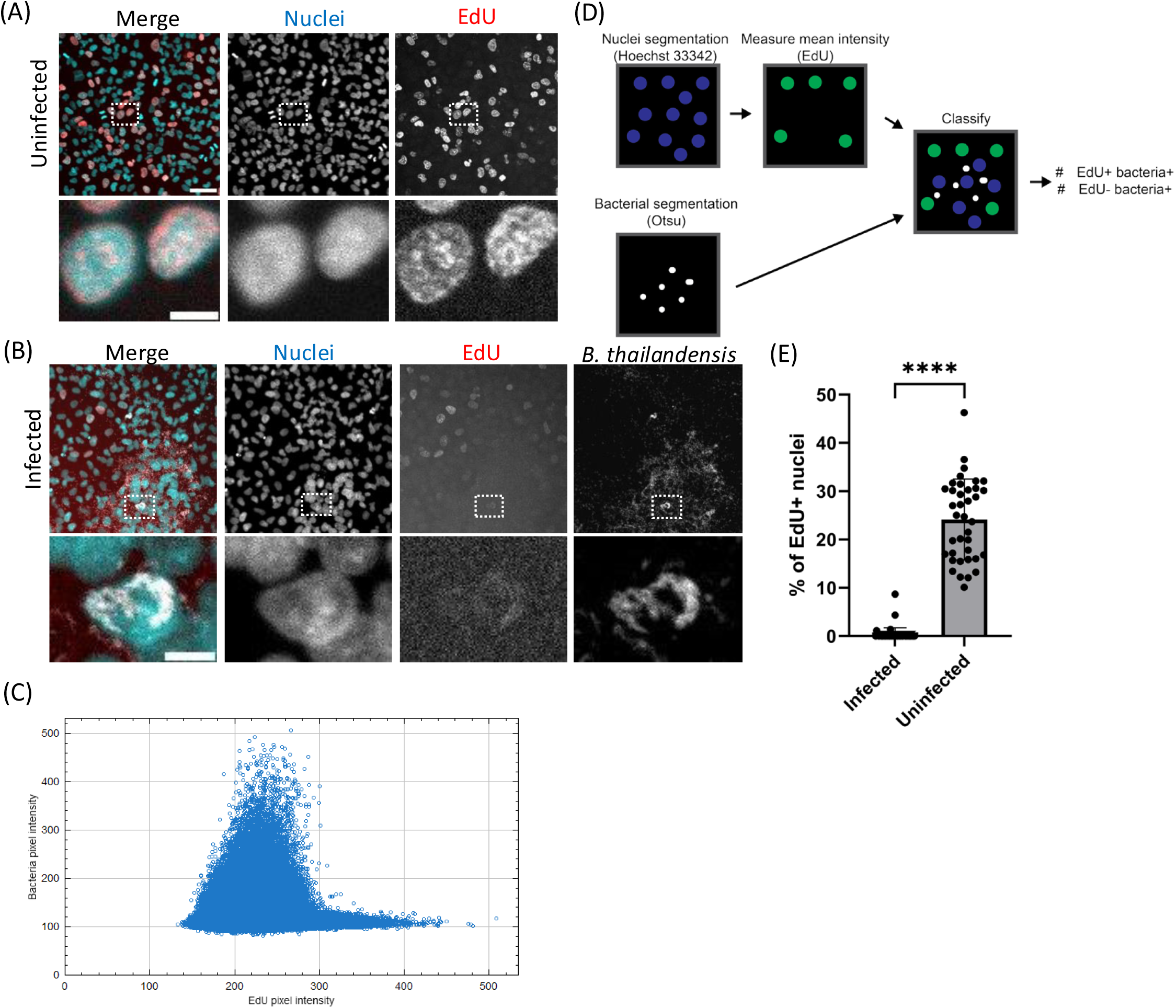
Infected cells and nuclei do not synthesize new DNA. **(A)** Example of uninfected A549 cells stained with Hoechst 33342 (cyan) and EdU (red). Scale bars represent 50 μm in the overview image and 10 μm in the magnified image. **(B)** Example of A549 cells infected with *B. thailandensis* at MOI 100. Infected cells were fixed at 15 hpi and stained with Hoechst 33342 (cyan) and EdU (red). Scale bar represents 10 μm. **(C)** Scatterplot of fluorescence intensity per pixel of the bacteria and EdU channel for the image shown in **(B)**. Bacterial pixel intensities were plotted for all pixels in the image, whereas EdU pixel intensities were plotted for pixels within segmented nuclei regions of interest (ROI). **(D)** Schematic representation of the image analysis workflow used to quantify EdU intensity and bacterial proximity. Bacteria were segmented using Otsu thresholding, and host nuclei were segmented from the Hoechst 33342 channel. The mean EdU fluorescence intensity was measured for each nuclear ROI. Nuclei were subsequently classified as EdU-positive or EdU-negative and as bacteria-proximal or bacteria-distal based on their association with segmented bacteria. **(E)** Quantification of the percentage of infected and uninfected nuclei positive for EdU incorporation. Each datapoint represents a single image. Data are from two independent biological replicates. An unpaired t-test with Welch’s corrections was carried out (**** P<0.0001).

To quantify overall DNA synthesis status of the infected monolayer, we used the Otsu algorithm to segment host nuclei and bacteria and then measured per-pixel signal intensity in the bacterial channel across the full image and in the EdU channel within segmented nuclear regions of interest (ROI). We plotted these values against each other and observed a negative correlation (Figure 3C). In addition, we also developed an image analysis pipeline to identify host cell nuclei positive for EdU with bacteria within 1 μm (Figure 3D, S3). In uninfected samples, 24% of nuclei incorporated EdU. In contrast, only 0.4% of nuclei in proximity to *B. thailandensis* incorporated EdU (Figure 3E). These results indicate that infected cells and nuclei rarely synthesize new DNA.

### Inhibition of cell division reduced the number of nuclear invasion events

When examining time-lapse images, we noticed that nuclear invasion events can happen after a prior mitotic event (Figure 4A). We therefore hypothesize that nuclear envelope breakdown during mitosis allows *B. thailandensis* to localize to future nuclear regions and become retained within daughter nuclei after division (Figure 4B). To determine whether mitosis is required for nuclear invasion, we inhibited mitosis using RO-3306 CDK inhibitor(44). A549 cells were incubated with RO-3306 for 8 hours prior to infection and were then infected as described above. Notably, nuclear invasion events were still observed in the presence of RO-3306 (Figure 4D). To quantify and compare the rate of nuclear invasion events between RO-3006-treated and DMSO-treated cells, we developed a semi-automated image analysis pipeline to segment and quantify infected cells. In brief, host cells were segmented using Cellpose(45), and bacteria were segmented using the Otsu algorithm. Mean fluorescence intensity in the bacterial channel was then used to define a threshold for classifying host cells as infected or non-infected (Figure 4C, S4A). To account for variation in infection rates between conditions, we normalized the number of nuclear invasion events to 1,000 infected cells. RO-3306 did not interfere with *B. thailandensis* infection as there was no significant difference in the percentage of infected cells (Figure S4B). However, there is a significant reduction in number of nuclear invasion events per 1000 infected cells in the RO-3306-treated cells (Figure 4E). On average, one invasion event was observed in 500-1000 DMSO-treated infected cells, compared to one in around 2000 RO-3306-treated cells. This indicates that most nuclear invasion events are the consequence of nuclear envelope breakdown during mitosis, however, additional mechanisms may play a role as well.

**Figure 4.**
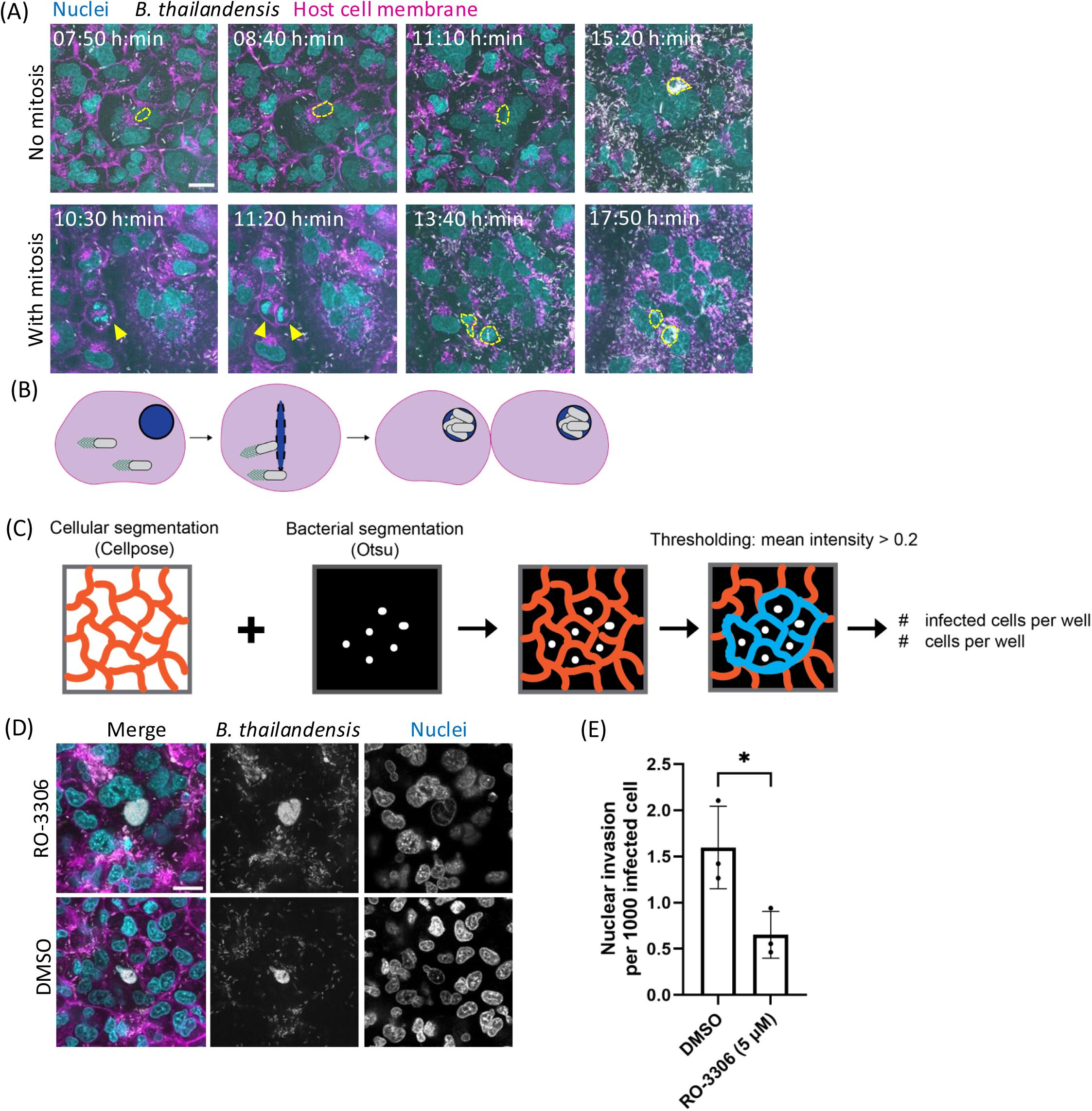
Mitosis facilitates bacterial entry into the nucleus, and its inhibition decreases nuclear entry rates. **(A)** Time-lapse images of nuclear invasion event with or without prior mitosis event. A549 cells were infected with *B. thailandensis* at MOI 100 and imaged from 5:30 hpi with 10-minute intervals. Host cell membrane was stained with CellMask DeepRed (magenta) and nuclei stained with Hoechst 33342 (cyan). Scale bar represents 20 µm. **(B)** Scheme model of the potential mechanism by which *B. thailandensis* enters host cell nuclei during mitosis. **(C)** Scheme depicting image analysis pipeline. Eukaryotic cells stained with CellMask DeepRed were segmented with Cellpose and fluorescent bacteria were segmented with the Otsu algorithm. The channels were then overlayed and the definition of an infected cell was threshold to a mean bacterial fluorescence of 0.2. **(D)** Representative images of *B. thailandensis* nuclear invasion events in A549 cells treated with either DMSO or RO-3306 (5 μM). Host cells were infected at MOI 100 and images were taken at 16 hpi. Host cell membrane was stained with CellMask DeepRed (magenta) and nuclei stained with Hoechst 33342 (cyan). Scale bar represents 20 µm. Images representative of at least 3 independent biological replicates. **(E)** Nuclear invasion events of *B. thailandensis* per 1000 infected cells in either DMSO-or RO-3306-treated cells. A549 cells infected with *B. thailandensis* at MOI 100 and images were taken with a 20x objective between 14 to 17 hpi. An unpaired Welch’s t-test was carried out (* P< 0.05).

### Actin tail motility is essential for nuclear invasion

We next hypothesized that nuclear invasion events could be dependent on virulence factors. We tested key virulence factors: the T6SS-5 (Δ*hcp-5*), type III secretion system (Δ*bsaM*), extracellular flagella (Δ*fliC1*), intracellular flagella (Δ*motA2*), or actin tail motility (Δ*bimA*). To compare the rate of nuclear invasion, a monolayer of A549 cells was infected at MOI 100 with the mutants and at least 20 fields of view (905.54 μm x 905.54 μm) were taken at late infection stages (14-16 hpi) for each mutant in each of the 3 biological replicates. Surprisingly, we discovered that all strains except for the Δ*bimA* mutant entered nuclei (Figure 5A, S5B). We observed that WT *B. thailandensis* invaded nuclei more often than the mutants (Figure S5C, left). However, since each mutant infects and spreads from cell-to-cell differently, we normalized the number of nuclear invasion events to the number of infected cells (Figure 5B, S5A). Importantly, this analysis only considered nuclear invasion events when the nucleus was filled with bacteria. Since bacteria can enter the nucleus at any point of infection, the actual number of nuclear invasion events could be higher. Nonetheless, with reference to our time-lapse images in Figure 1, the infection timepoint we chose for quantification (14-17 hpi) is a timepoint when nuclear invasion events are most apparent and there is minimal amount of death of the infected cells.

**Figure 5.**
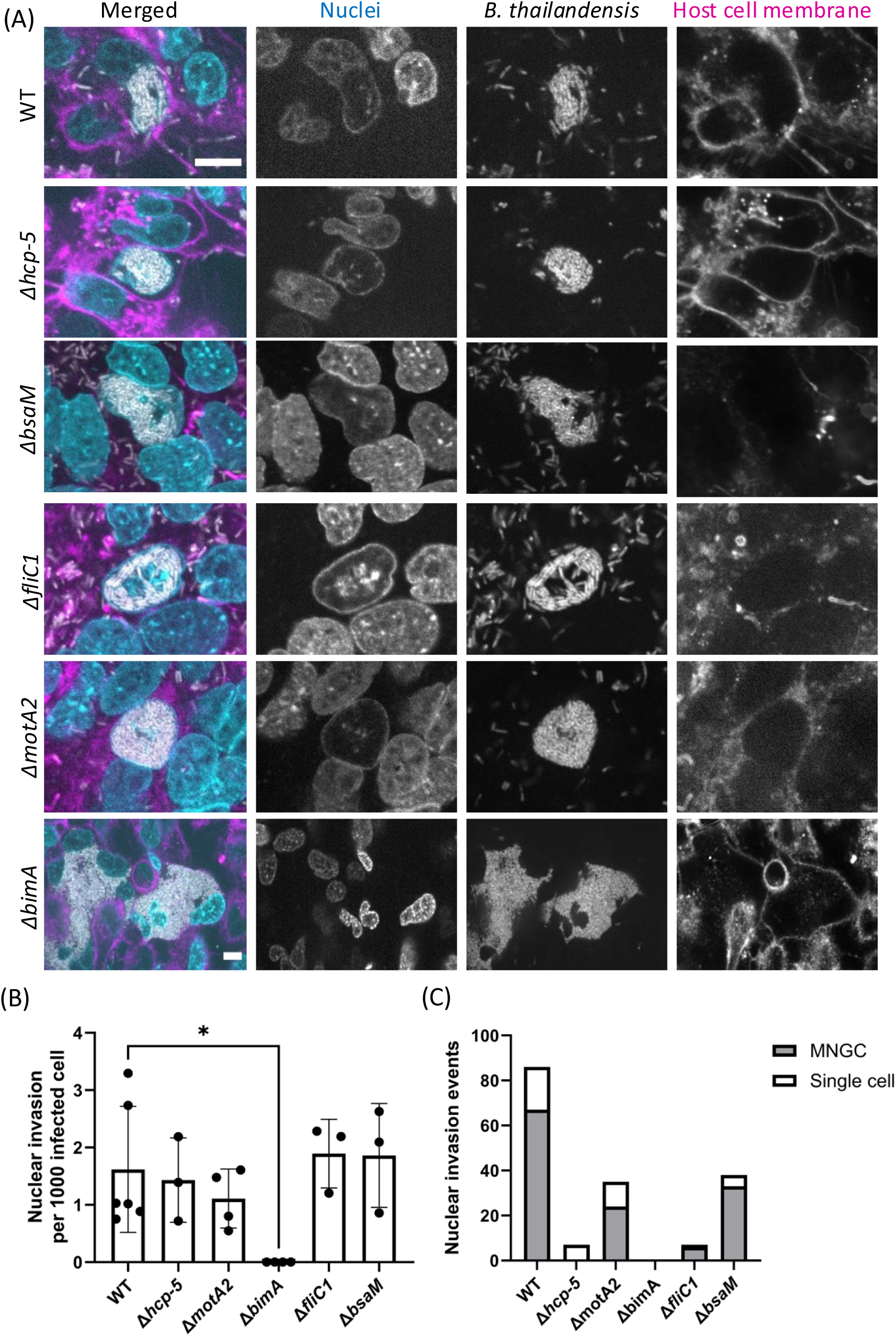
Actin tail motility is required for *B. thailandensis* to enter host cell nuclei. **(A)** Images of A549 nuclei invaded by various mutants of *B. thailandensis*. The cells were infected at MOI 100 and images were taken between 14 to 17 hpi. Host cell membrane was stained with CellMask DeepRed (magenta) and nuclei stained with Hoechst 33342 (cyan). Scale bar represents 10 µm. **(B)** Nuclear invasion events of various strains per 1000 infected cells. A549 cells infected with various mutants of *B. thailandensis* at MOI 100 and images were taken with a 20x objective between 14 to 17 hpi. One-way ANOVA with Tukey’s multiple-comparisons test was used to determine the statistical significance (ns: not significant, * p<0.05, ** p<0.01, *** p< 0.001, **** p< 0.0001). **(C)** Distribution of nuclear invasion events in MNGCs and single cells across different *B. thailandensis* mutant strains. The nuclear invasion events from the same dataset from **(B)** were manually assessed and classified as occurring within an “MNGC” or in a “single cell”.

Using the automated image analysis, we show that *B. thailandensis* WT infected 44% of the total number of cells, the T6SS-5 mutant (Δ*hcp-5*) infected 10%, Δ*motA2* mutant infected 38%, and Δ*fliC1* mutant infected 21%. The Δ*bsaM* mutant has a delayed phagosomal escape and only infected 0.7% of total number of cells. Moreover, since actin tail motility is important for cell-to-cell spread, the Δ*bimA* mutant only infected 1% of the monolayer (Figure S5C, right). After normalization of nuclear invasion events to the number of infected cells, we found that amongst 70,331 cells infected with WT, 86 nuclei contained intranuclear bacteria, with 67 of these nuclei residing in MNGCs and 19 in single cells. For Δ*hcp-5* mutant, in 6,161 infected cells, we found 7 nuclear invasion events in single cells. For Δ*fliC1* mutant, in 21,180 infected cells, we found 33 invasion events in MNGCs and 5 in single cells. For Δ*motA2* mutant, in 33,504 infected cells, we found 24 invasion events in MNGCs and 11 in single cells. For Δ*bsaM* mutant, in 3,708 infected cells, we found 6 invasion events in MNGCs and 1 in a single cell. Overall, a nuclear invasion event could be found in 500-1000 cells infected either with WT, Δ*hcp-5*, Δ*fliC*, or Δ*motA2*, or Δ*bsaM* (Figure 5B), and the majority of nuclear invasion events were identified in MNGCs (Figure 5C). For Δ*bimA* mutant, we analyzed 14,945 infected cells, therefore expected about 15-30 unclear invasion events if the rate of nuclear invasion was independent of *bimA*, however, we observed no nuclear invasion events. This suggests that actin-based motility is critical for nuclear invasion.

### Nuclear invasion is independent of the mechanism of actin nucleation

To test if another intracellular pathogen that forms actin tails can also invade nuclei, we infected A549 cells with *Shigella flexneri*. *S. flexneri* express IcsA, which recruits the host N-WASP complex for forming actin tails(46–48). We observed 40,655 A549 cells infected with *S. flexneri* expressing dsRed between 14 to 16 hpi and counted 21 nuclear invasion events (Figure 6A, C). This suggests that although actin tail is essential for nuclear invasion, the mechanism of actin nucleation has no effect. In addition, nuclear invasion events are rarer for *S. flexneri*, occurring 1 in 2,000 infected cells (Figure 6C). In *S. flexneri*, IcsA was shown to be required for host cell invasion and the *icsA* mutant cannot form actin tails to spread from cell-to-cell(46,49), therefore, testing if actin tail formation is required for nuclear invasion using *icsA* mutant is confounded by 100-fold lower number of infected cells. Nevertheless, we infected A549 cells with a *S. flexneri* strain lacking IcsA and observed no nuclear invasion events in 5,565 infected cells (Figure 6B,C) suggesting that indeed actin tail formation is also required for nuclear invasion by *S. flexneri*.

**Figure 6.**
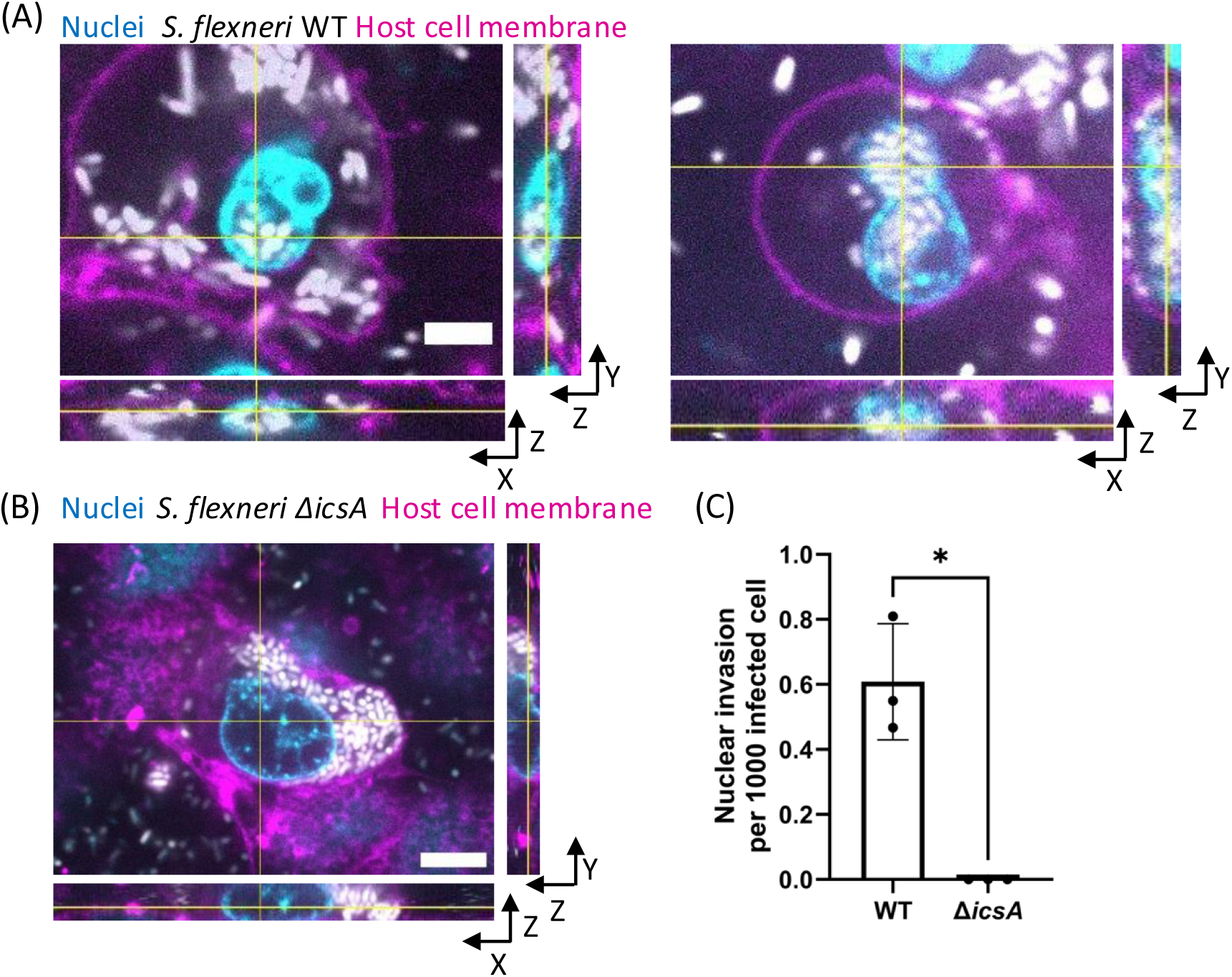
*S. flexneri* invades A549 cell nuclei by actin tail motility. **(A)** Z-stack images of intranuclear *Shigella flexneri.* A549 lung cells were infected with *S. flexneri* expressing ds-Red (grey) on the pMW211 plasmid at MOI of 10. Infection was performed in Opti-MEM^TM^ supplemented with 30 μg/mL of gentamicin. Images were taken at 15 hpi. Host cell membrane was stained with CellMask DeepRed (magenta) and nuclei stained with Hoechst 33342 (cyan). Orthogonal views covering the whole 25 μm z-stack are displayed. Scale bar represents 5 µm. **(B)** Z-stack images of cytosolic *Shigella flexneri* Δ*icsA.* A549 lung cells were infected with *S. flexneri* Δ*icsA* expressing ds-Red (grey) on the pMW211 plasmid at MOI of 500. Images were taken at 16 hpi. Host cell membrane was stainedm with CellMask DeepRed (magenta) and nuclei stained with Hoechst 33342 (cyan). Orthogonal views covering the whole 10.5 μm z-stack are displayed. Scale bar represents 10 µm. **(C)** Nuclear invasion events per 1000 cells infected with *S. flexneri* WT or Δ*icsA*. A Welch’s unpaired t-test was performed (n=3, * P<0.05).

### Inhibition of actin polymerization prevents nuclear invasion

To further test the role of actin motility in nuclear invasion by *B. thailandensis,* we inhibited actin polymerization by cytochalasin D, which was used in studies to block actin motility of intracellular bacteria(50,51). We infected A549 cells with *B. thailandensis* WT for 4 hours to allow for invasion and phagosomal escape, then treated the infected cells with either 5 µg/mL of cytochalasin D or 0.5% of DMSO. Representative images showed nuclear invasion events in DMSO-treated cells, whereas cytochalasin D-treated cells contained intracellular bacteria but no detectable nuclear invasion events (Figure 7A–C). We then applied our image analysis pipeline to images acquired at the final time point (15–17 hpi) and quantified nuclear invasion events normalized to the number of infected cells. On average, nuclear invasion events were detected 1 in 250 DMSO-treated cells infected with *B. thailandensis*, whereas no nuclear invasion events were detected in 2,664 cytochalasin D-treated cells infected with *B. thailandensis* (Figure 7D). These results support the conclusion that actin polymerization is important for nuclear invasion.

**Figure 7.**
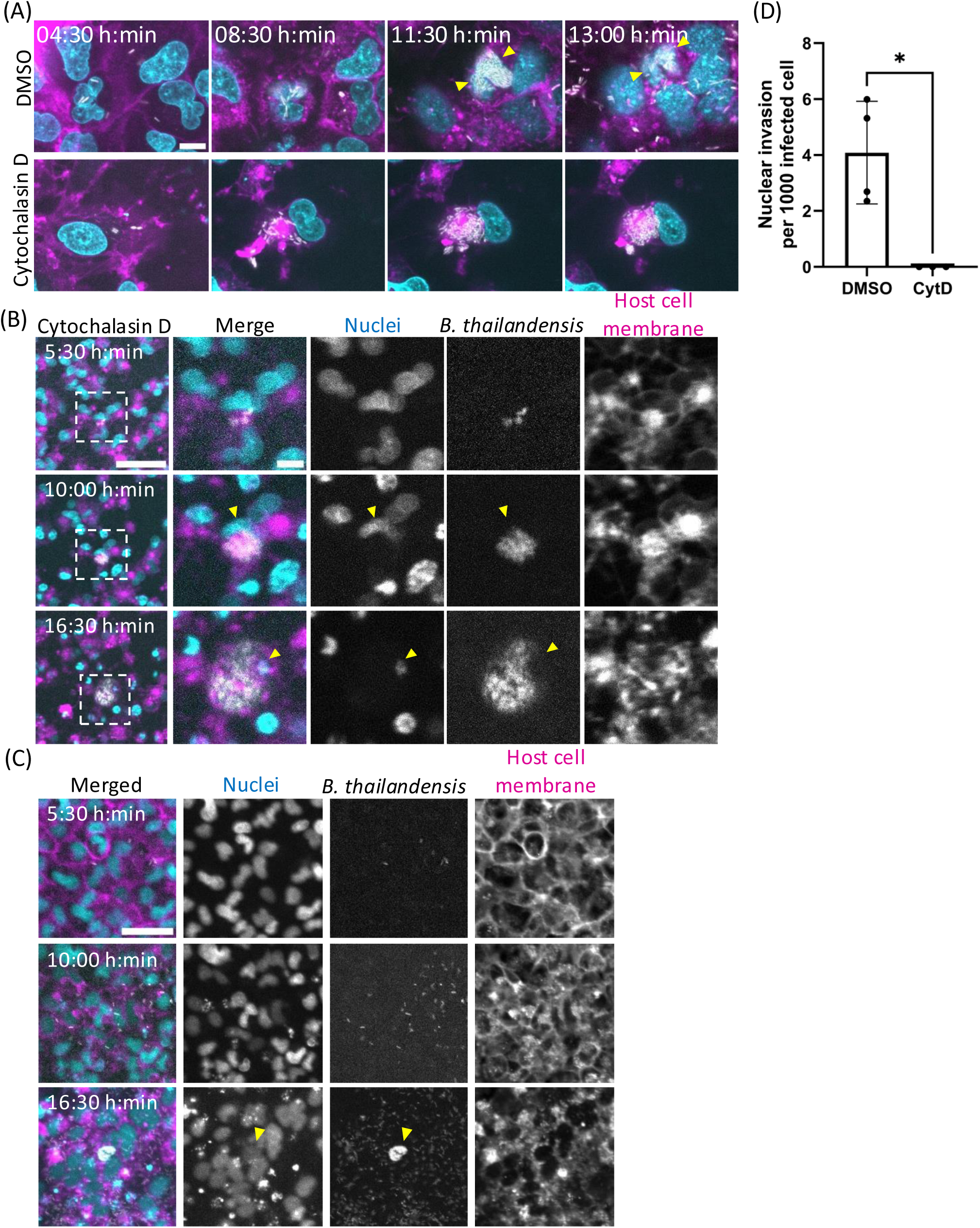
Cytochalasin D treatment abolishes nuclear invasion events. **(A)** Time-lapse images of A549 cells infected with *B. thailandensis* expressing TssB-5-mScarlet-I (grey) at MOI of 50, and treated with cytochalasin D (5 µg/mL) or 0.05% of DMSO from 4 hpi. Images were taken at 15-minute intervals. Host cell membrane was stained with CellMask DeepRed (magenta) and nuclei stained with Hoechst 33342 (cyan). Yellow arrows indicate nuclear invasion events in the cells treated with DMSO. Scale bar represents 10 µm. **(B,C)** Time-lapse images images of A549 cells infected with *B. thailandensis* expressing TssB-5-mScarlet-I (grey) at MOI of 50, and treated with **(B)** cytochalasin D (5 µg/mL) or **(C)** DMSO at 4 hpi. Images were taken with a 20x objective and at 30 min intervals. Host cell membrane was stained with CellMask DeepRed (magenta) and nuclei stained with Hoechst 33342 (cyan). Yellow arrows highlight nuclei of the infected cell. Scale bars represent 50 µm in the overview image and 10 µm in the magnified image. **(D)** Nuclear invasion events per 1000 infected cells treated with DMSO or cytochalasin D. A Welch’s unpaired t-test was performed (n=3, * P<0.05).

## Discussion

In this study, we show that *B. thailandensis* invades host cell nuclei and replicates intranuclearly. This process is mostly promoted by host cell mitosis. However, by generating in-frame deletions of various virulence factors and chemically inhibiting actin polymerization, we identify that actin tail motility is essential for *B. thailandensis* to enter host cell nucleus.

Our work mostly focused on the A549 epithelial lung cell line, but we also observed nuclear invasions in HeLa cells (Figure 1D). Since it was also reported that *B. pseudomallei* localized to the nucleus of infected human lung and guinea pig spleen tissue(52), it is likely that *Burkholderia* invasion to the nucleus occurs in a wide range of cells *in vitro* as well as *in vivo*.

We noticed that nuclear invasion events occurred more frequently within MNGCs (Figure 5C). This enrichment is likely because most infected cells are contained within MNGCs (see overview images in Figure S5B). Given the incidence of a nuclear invasion event is 1 in 500-1,000 infected cells, it is statistically more likely for such rare events to be detected in cells residing within MNGCs. Although MNGCs are formed by T6SS-5 activity, it is important to note that the T6SS-5 deletion mutant can also invade host nuclei, indicating that nuclear invasion events are not dependent on T6SS-5.

Given that the nuclear compartment is typically shielded by the nuclear envelope, we examined whether *B. thailandensis* could exploit nuclear envelope breakdown during mitosis to gain nuclear access. Our findings suggest that nuclear envelope breakdown during mitosis represents one route by which *B. thailandensis* gains access to the nucleus, as mitotic events often occurred prior to the detection of intranuclear bacteria (Figure 4A). However, since treatment with RO-3306 did not completely eliminate nuclear invasion events, mitosis may not be the sole mechanism.

We found that virulence factors such as the T3SS, T6SS-5 and flagella have no effect on the rate of nuclear invasion events but actin tail formation is essential (Figure 5A,B). Furthermore, such nuclear invasion events are also observed for *Shigella flexneri* (Figure 6). No nuclear invasion was observed in *B. thailandensis* and *S. flexneri* strains unable to polymerize actin (Figure 5A and 6C). Another bacterium that enters host nuclei, *Rickettsia* spp., forms actin tails with RickA and Sca2, which utilizes different host actin nucleators at different stages of infection(53–55). This suggests that although actin tail formation is essential for entering the host nucleus, the mechanism of actin nucleation is inconsequential. However, the precise mechanism by which the bacterium uses actin tails to enter the nucleus is unclear. One possibility is that nuclear entry occurs through the direct application of mechanical force, with actin tail motility generating sufficient force to penetrate the nuclear envelope, whereas the force generated by flagella is insufficient. Another possibility is that *B. thailandensis* exploits transient openings in the nuclear envelope. Transient nuclear envelope rupture events have been reported in the interphase cells and can result in a temporary loss of nuclear compartmentalization(56). In this model, actin tail-driven propulsion could provide the force necessary for the bacterium to pass through these openings.

Additionally, we tested the role of actin tail motility in nuclear invasion by chemically inhibiting host actin polymerisation with cytochalasin D(50,51). As actin is a core component of the cytoskeletal network that maintains cellular architecture, cytochalasin D treatment caused host cells to appear structurally weakened and less rigid. We allowed a 4-hour infection period for the bacteria to reach the host cell cytoplasm before adding cytochalasin D. After treatment, WT *B. thailandensis* was unable to form actin tails and remained confined to a single infected cell, resulting in an infection phenotype similar to that observed for the Δ*bimA* mutant. The absence of nuclear invasion events under cytochalasin D treatment further supports the critical role of actin-based motility in this process (Figure 7).

*Candidatus Nucleicultrix amoebiphila* was shown to enter host cell nuclei during cell division when the nuclear envelope dissolves(11). However, *B. thailandensis*, at least *in vitro*, seems to also depend on actin polymerization. Indeed, the non-motile *bimA* mutant, which replicates within the host cells to high density, fails to invade the nucleus. Since actin tail formation is essential for *B. thailandensis* to enter host cell nuclei, we speculate that they enter host nucleus by mechanical force generated by actin tail motility. However, it is unlikely that *B. thailandensis* enter host cell nuclei via the nuclear pore complex since the pore size is too small to allow a bacterium to pass through (∼50 nm)(57,58). It is also possible that *B. thailandensis* secrete effectors to facilitate nuclear entry. *Horospora* spp. express a 89 kDa protein that allows them to bind to the nuclear membrane and facilitate entry(59).

Interestingly, we never observed any intranuclear *B. thailandensis* leaving the host nucleus. The presence of actin inside of the host nucleus have been reported(60–63), but the form of actin inside of the nucleus is unclear(64–66). It is possible that *B. thailandensis* is unable to initiate actin polymerization inside of the nucleus and thus cannot form actin tails to leave of the nucleus. This is also supported by the observations that the intranuclear bacteria are mostly non-motile (Video S1).

Replicating inside of the host nucleus has certain potential advantages for the pathogen. The nuclear niche could provide protection against innate host defense such as autophagy. Moreover, the intranuclear environment is nutritious. Symbiotic intranuclear bacteria such as *Rickettsiae* spp. and *Chlamydiae* are known to feed on host nucleotides by ATP/ADP translocases(67,68). In fact, a reduction of host heterochromatin has been observed for intranuclear bacteria(69). Curiously, we also observed a reduction in Hoechst 33342 signal in nuclei containing *B. thailandensis*, however it is unclear if this was due to lower DNA content or altered Hoechst fluorescence due to local acidification associated with bacterial accumulation (Figure 3,5A).

Intranuclear bacteria are mostly studied as symbionts in amoeba and ciliates and are rarely studied in the context of mammalian cell infection(11,70). The role of intranuclear bacteria in latent and relapsing infections are yet to be explored. It is possible that the intranuclear environment can act as a protective niche and allow for secondary infections when dead cells containing intranuclear bacteria are phagocytosed by immune cells like macrophages. For example, *Francisella tularensis* replicates in a vacuole that can be trogocytosed, leading to a secondary infection in phagocytes(71,72). Furthermore, certain cell types, including intestinal epithelial cells and skin keratinocytes, undergo frequent mitosis(73,74). Moreover, alveolar epithelium cells in the lung also regenerates and divide after lung injury(75). Since the gut, skin, and lung are considered the primary infection sites for *Burkholderia pseudomallei*(16,76), the transient remodeling of the nuclear envelope during cell division in these cells may provide opportunities for pathogens to access the nucleus.

In summary, we show that *B. thailandensis* and *S. flexneri* can gain entry into the nucleus during mitosis, and may also do so under non-mitotic conditions by utilizing actin tail motility. This work expands current knowledge on intracellular bacterial niches and enables further research into mechanisms of entry to the intranuclear niche, especially given that mitotically active cell populations are present in many human tissues.

## Materials and Methods

### Mammalian cell lines and culturing

A549 lung cells (CCL-185, ATCC) were cultured in Ham’s F-12 media (Sigma-Aldrich). HeLa cells (CCL-2, ATCC) were cultured in DMEM (Sigma-Aldrich). All of the culturing media used was supplemented with 10% heat-inactivated fetal calf serum (FCS, BioConcept AG). The cell lines used were cultured at 37°C with 5% CO2 and used for experiments from passages 2 to 10.

### Construction of genetic mutants

Genetic deletion mutants were constructed via allelic exchange via the vector pDONRPEX-18Tp-SceI-pheS(77). The homologous flanking regions were at least 1000 bp. The assembled plasmids were conjugated into *B. thailandensis* strains by *E. coli* SM10 λpir. Positive conjugants were selected by trimethoprim resistance and counter selection were performed on M9 minimal medium agar plates supplemented with 0.4% glucose and 0.1% (w/v) 4-chloro-phenylalanine. Mutants were verified by polymerase chain reaction (PCR) and Sanger sequencing. Constitutive expression of proteins were facilitated by the plasmid pUC18T-mini-Tn7-Tp and helper plasmid pTNS2 under the ribosomal promoter pSC12. This inserts the fragments at the *attn7* stie near the *glmS* locus(78). Bacterial strains are listed in Table 1.

**Table 1.**
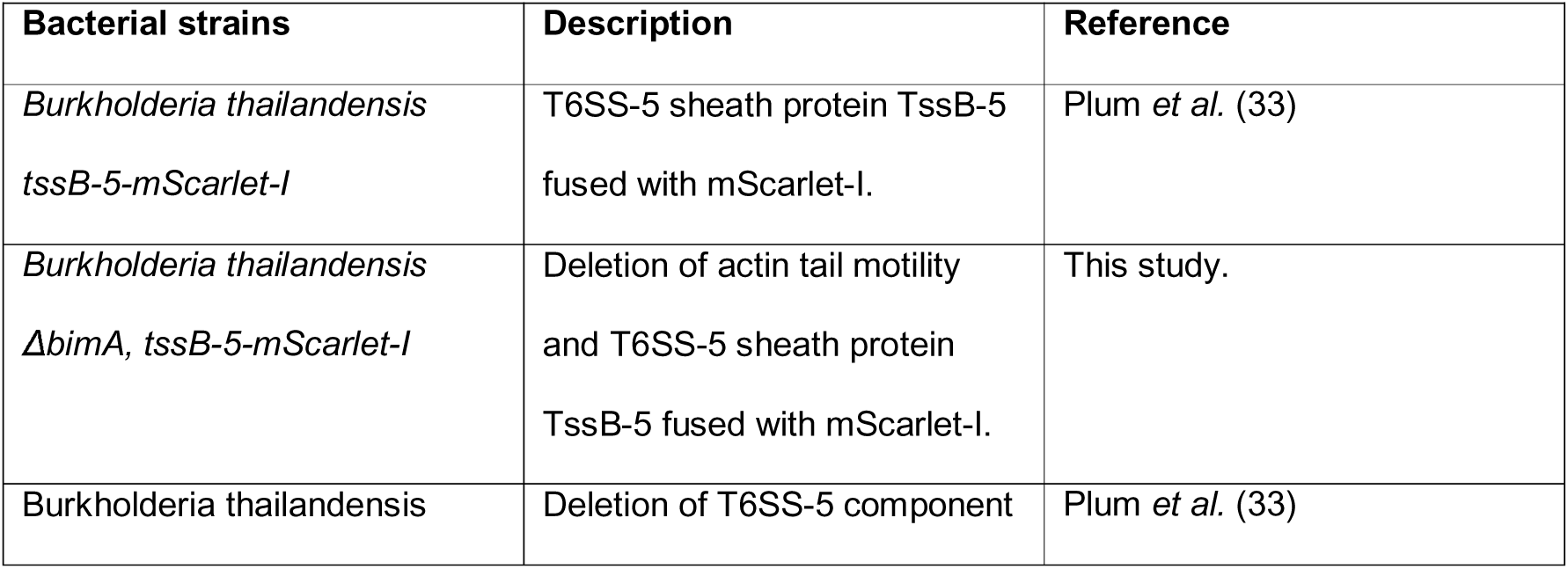
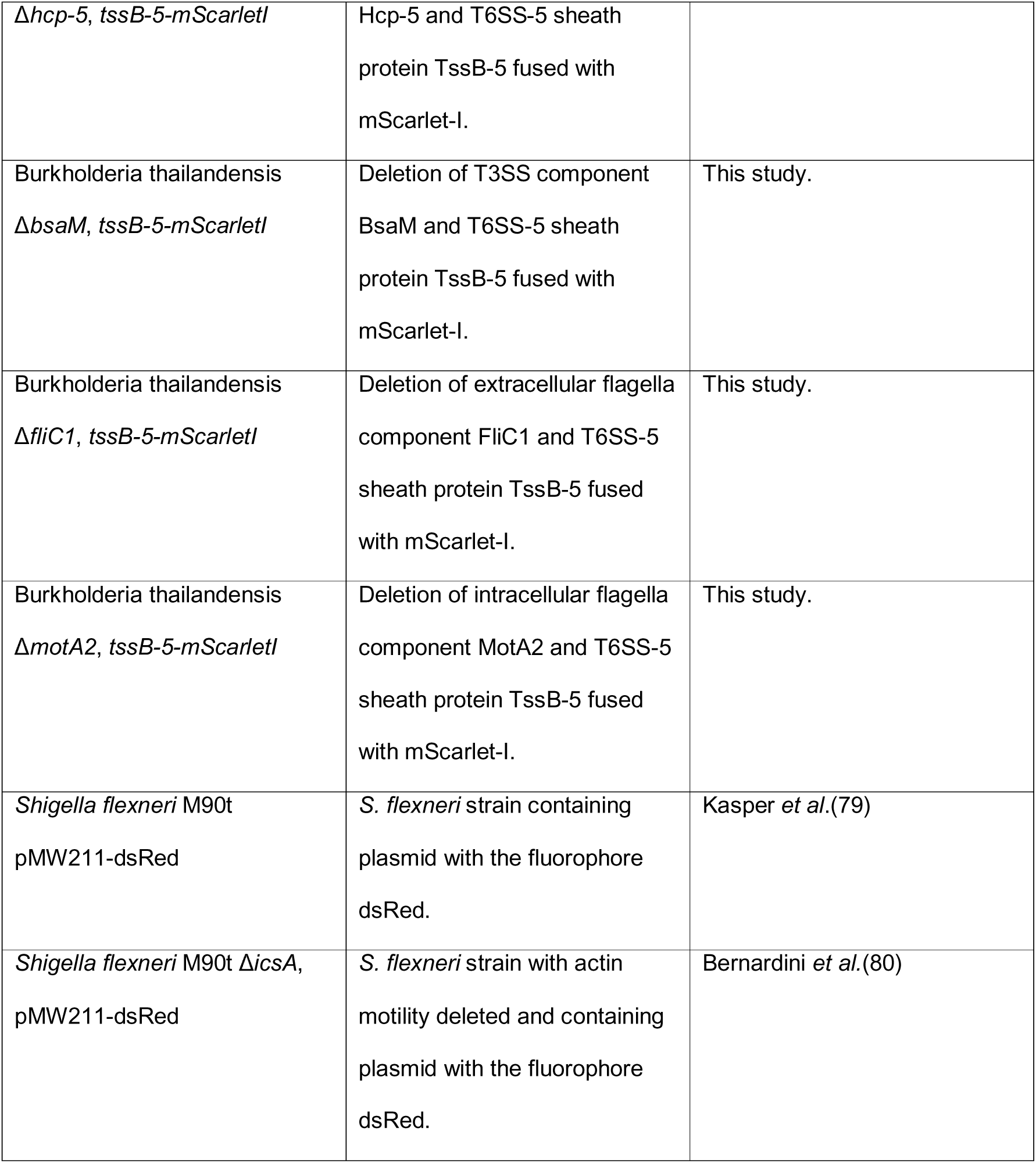
List of bacterial strains used in this study.

### Infection of host cells

A549 lung cells and HeLa cells were seeded at 8 x 10^4^ cells per well on µ-slide 8 well glass bottom (Ibidi GmbH) 24 hours before infection. The bacterial strains used for infection were grown overnight at 37°C at 200 rpm in low salt LB. Prior to infection, the bacterial cultures were diluted (1:100) and grown until mid-log phase (OD 0.4-0.8). Host cells and bacteria were washed twice in Opti-MEM^TM^ Reduced Serum Medium (Sigma-Aldrich). The bacteria were then added to the cells at MOIs specified for every experiment and the slides were centrifuged at 300 x g for 5 min to synchronize infection. After 1 hour post infection (hpi), the cells were washed with Opti-MEM^TM^ supplemented with either 300 μg/mL of kanamycin (for *B. thailandensis*) or 30 μg/mL of gentamicin (for *S. flexneri*).

### Inhibition of mitosis

Eukaryotic cells were seeded as described above. Mitosis inhibitor RO-3306 (5 μM, Sigma Aldrich) was added 8 hours in culture medium prior to infection and was replenished in Opti-MEM^TM^ Reduced Serum Medium after cells were infected. The inhibitor was maintained in the culture medium throughout the experiment and replenished after cell staining prior to imaging.

### Inhibition of actin polymerization

Cells were infected as described above. Cytochalasin D stock solution in DMSO (10 mg/mL, Sigma Aldrich) was diluted to a final concentration of 5 µg/mL in Opti-MEM^TM^ Reduced Serum Medium containing either 300 μg/mL of kanamycin or 30 μg/mL of gentamicin. The diluted solution was added to the infected cells at 4 hpi. To control for the effects of the solvent, 1.5 µL of DMSO was added to a separate well of infected cells containing 300 µL of Opti-MEM^TM^ to reach 0.05%.

### Detection of DNA synthesis

EdU incorporation and staining were performed according to the manufacturer’s protocol (ThermoFischer). Briefly, cells were seeded one day prior to infection and either infected as described above or left uninfected. At the indicated time points, EdU was added to a final concentration of 20 μM, and the cells were incubated for 20 min. The EdU-containing medium was then removed, and the cells were fixed with 3.7% paraformaldehyde (PFA) for 15 min at room temperature. Following fixation, cells were washed twice with 2% bovine serum albumin (BSA) and permeabilized with 0.5% Triton X-100 for 20 min. After two additional washes with 2% BSA, the Click-iT reaction cocktail was added, and the cells were incubated for 30 min according to the manufacturer’s instructions. Host cell nuclei were subsequently stained with 40 μM Hoechst 33342 for 5 min, followed by two washes with 3% BSA before imaging.

### Confocal imaging

Images were recorded with Kinetix sCMOS camera using a 60x oil immersion objective (NA = 1.42; field of view – 302.28 x 302.28μm) and 20x air objective (NA = 0.8; field of view – 905.54 μm x 905.54 μm) on the Nikon ECLIPSE Ti2 with the Crestoptics X-Light V3 spinning disk confocal system at 37°C with 5% CO_2_. To visualize eukaryotic plasma membrane, cells were stained with CellMask DeepRed (ThermoFischer) at a concentration of 5 µg/mL for 5 min at 37°C with 5% CO_2_. To visualize host cell nuclei, the cells were stained with Hoechst 33342 at 40 µM for 5 min at 37°C with 5% CO_2_.

### Quantification of EdU fluorescence intensity

Multichannel fluorescence images were converted into two-dimensional maximum-intensity projections across all z-planes prior to analysis. Host cell nuclei were identified from the Hoechst 33342 channel. Images were smoothed using a Gaussian filter (σ = 2 pixels) and thresholded using Otsu’s method to generate binary masks. Holes within segmented nuclei were filled and touching nuclei were separated using watershed segmentation. Nuclear regions of interests (ROI) were identified using the Analyze Particles function in ImageJ.

To identify bacteria, the bacterial fluorescence channel was subjected to background subtraction by subtracting a Gaussian-blurred image (σ = 20 pixels) from the original image. The resulting image was thresholded using Otsu’s method and converted into a binary mask.

For each nucleus, mean EdU fluorescence intensity was measured within the nuclear ROI. Nuclei with mean EdU intensity above 270 a.u. were classified as EdU-positive. To assess bacterial association, each nuclear ROI was expanded by 1 μm and the mean intensity of the bacterial mask within the expanded region was measured. Cells were classified as bacteria-positive when the mean fluorescence intensity is above 10 a.u.. Based on these criteria, cells were categorized as EdU-positive, bacteria-positive, double-positive, or negative for both markers. For each image, the total number of nuclei, EdU-positive nuclei, bacteria-associated nuclei, and EdU-positive bacteria-associated nuclei were quantified. Segmentation masks and ROI annotations were visually inspected and exported for quality control.

For pixel-wise fluorescence intensity analysis, nuclei were segmented as mentioned above. Pixel intensities from the EdU and bacterial fluorescence channel were extracted on a pixel-by-pixel basis. Pixels contained in the nuclear ROIs were included for the Edu channel to exclude background regions. Global pixel counts for the bacteria channel was included. Scatter plots were generated by plotting EdU fluorescence intensity against bacterial fluorescence intensity for all nuclear pixels. For each image, segmentation masks and scatter plots were saved for quality control and visualization.

### Quantification of bacterio-nuclear colocalization per infected cells

A549 lung cells were seeded, infected and stained with CellMask DeepRed and Hoechst 33342 as previously described. The wells were imaged with a 20x objective between 14 to 16 hpi. At least 20 fields of view (FOV) were imaged per well.

In order to determine the number of infected cells, we developed an image analysis pipeline (Script available at: https://doi.org/10.5281/zenodo.10581517). Cells were segmented with Cellpose, a deep-learning cellular segmentation method to establish cell shapes(45). The cytoplasm model 2.0 was used for the segmentation. It was trained on 2-channel images – a membrane or cytoplasm and a nuclear channel with over 70’000 segmented objects. Every image was loaded from the OMERO interface on the Cellpose environment, where the cellular size was approximated to 40 µm and the plasma membrane channel was used as base for the segmentation. The nuclear channel was additionally used for the segmentation to get better spacing between cells in the dense monolayer. The segmented cells were saved as regions of interest (ROIs). Bacteria were segmented based on the Otsu algorithm, where pixels are categorized into foreground and background depending on the fluorescence intensity of the bacterial channel. The saved ROIs were combined with the segmented bacteria. The average bacterial fluorescence per ROI was measured. The threshold for classifying a cell as infected was set at a mean bacterial fluorescence of 0.2 for *B. thailandensis* and 10 for *S. flexneri*, reflecting differences in fluorophore brightness. This approach resulted in 2 classes of segmented cells – infected and non-infected. Cellular and bacterial segmentation was manually evaluated by overlaying the membrane and bacterial channel to ensure a reliable segmentation.

Colocalization of bacteria and host nuclei was assessed manually. The number of colocalization events was then normalized to the number of infected cells.

### Immunofluorescence

To visualize *B. thailandensis* actin tail formation, phalloidin staining was performed on infected A549 cells. Cells were washed twice with PBS at 10-12 hpi and fixed with 4% methanol-free paraformaldehyde for 15 min at room temperature. The sample was washed twice and permeabilized with 0.1% Triton^TM^ X-100 for 15 min and subsequently washed twice again. The fluorescent Phalloidin (Alexa Fluor™ Plus 647 Phalloidin, Thermo Fisher) staining solution was prepared at a final concentration of 12.5 µg/ml in PBS supplemented with 1% BSA. The cells were stained with Phalloidin for 30 to 60 min at room temperature and were then stained with Hoechst 33342 as described previously. Afterwards, the cells were washed twice before imaging.

## Acknowledgments

The authors would like to thank Laurent Guerard, from the Imaging Core Facility of the Biozentrum for his assistance in developing our image analysis pipeline, the Dehio group for providing the *Shigella flexneri* WT strain, Prof. Dr. Philippe Sansonetti for providing the *Shigella flexneri* Δ*icsA* strain. This work was supported by the Boehringer Ingelheim Fonds PhD fellowship, the University of Basel, and the European Research Council, consolidator grant 865105 – “AimingT6SS”.

## Author contributions

Conceptualization, H.C.C., P.R.I., M.T.W.P., and M.B.; Methodology, H.C.C. and P.R.I.; Investigation, H.C.C. and P.R.I.; Writing – original draft, H.C.C.; writing – review and editing, H.C.C., P.R.I., and M.B.; funding acquisition, M.B.; supervision, M.B.

## Supplementary figure legends

**Figure S1.**
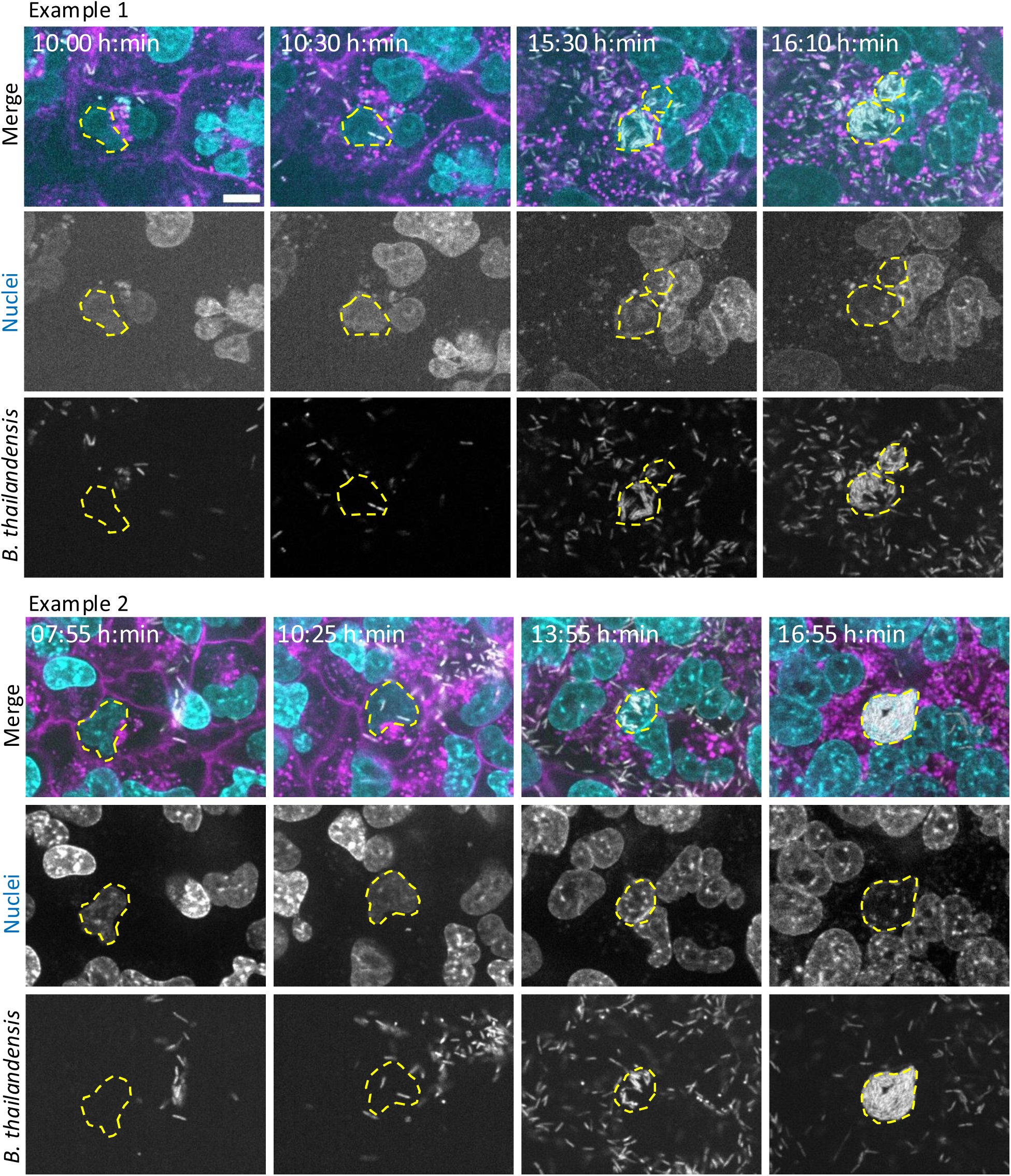
*B. thailandensis* invades A549 cell nuclei. Two representative examples of nuclear invasion events captured by time-lapse imaging. A549 lung cells infected with *B. thailandensis* expressing TssB-5-mScarlet-I (grey) at MOI of 50. Infection was performed in Opti-MEM^TM^ supplemented with 300 μg/mL of kanamycin. Host cell membrane was stained with CellMask DeepRed (magenta) and nuclei stained with Hoechst 33342 (cyan). Nuclei containing *B. thailandensis* are outlined with yellow dotted lines. Scale bar represents 10 µm.

**Figure S2.**
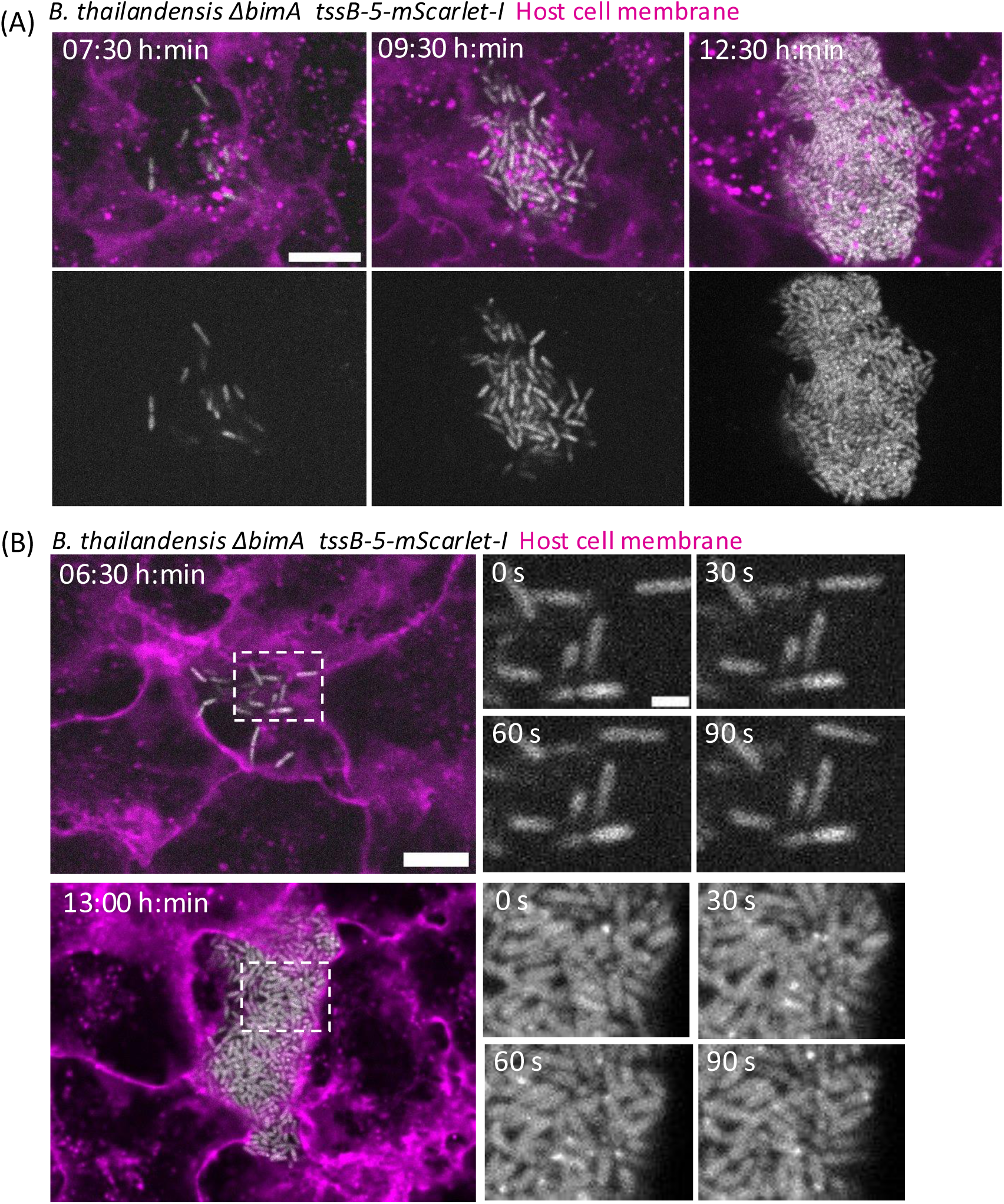
T6SS-5 assemblies of *B. thailandensis* Δ*bimA* mutant. **(A)** Time-lapse images of A549 cells infected with *B. thailandensis* Δ*bimA* expressing TssB-5-mScarlet-I (grey) at MOI of 50. Infection was performed in Opti-MEM^TM^ supplemented with 300 μg/mL of kanamycin. Images were taken at 10-minute intervals. Host cell membrane was stained with CellMask DeepRed (magenta). Scale bar represents 10 µm. **(B)** T6SS-5 assemblies observed in cytosolic *B. thailandensis* Δ*bimA*. A549 cells were infected with *B. thailandensis* Δ*bimA* expressing TssB-5-mScarlet-I (grey) at MOI of 50. Confocal images were taken at 30 second intervals for a duration of 15 minutes at 6:30 hpi and 13 hpi. Scale bar on the larger image represents 10 μm and 2 μm in the magnified images.

**Figure S3.**
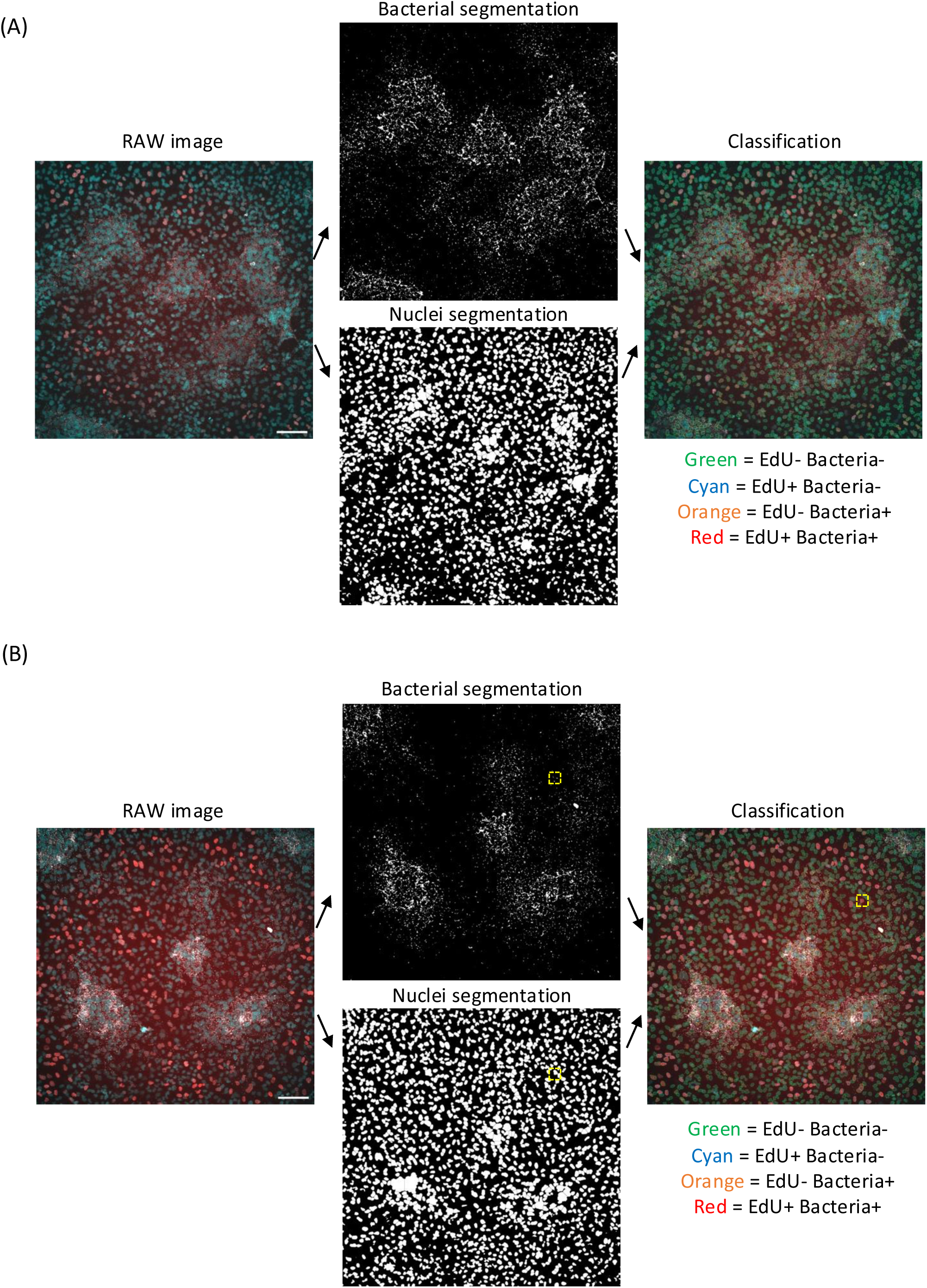
Examples of segmentation and classification of infected cells with or without DNA synthesis. **(A)** Raw image, segmented bacteria, and segmented nuclei are shown. Nuclei were classified into four categories and color-coded accordingly. **(B)** Same classification pipeline as in **(A)**, with an example of an EdU-positive, bacteria-positive nucleus highlighted by the yellow dashed box.

**Figure S4.**
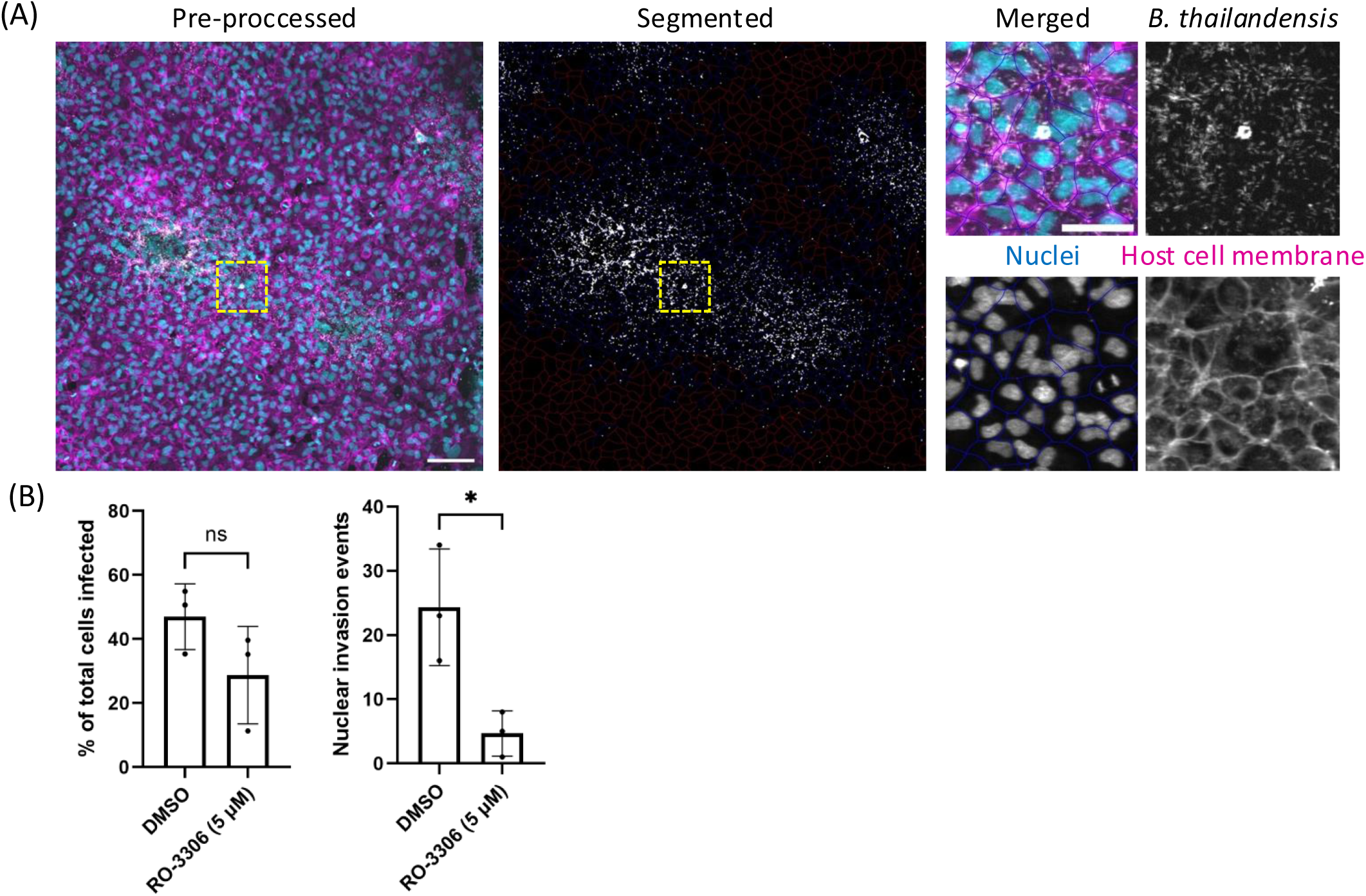
Segmentation of infected cells and quantification of nuclear invasion rates under conditions of mitotic inhibition. **(A)** Pre-processed image showing nuclei in cyan, bacteria in gray, and host cell membranes in magenta, alongside the corresponding segmented bacteria image. A magnified view of the region outlined by the yellow dashed box illustrates host cell segmentation using Cellpose. Scale bars represent 100 μm in the overview image and 50 μm in the magnified image. **(B)** Quantification of nuclear invasion events and the percentage of cells infected by *B. thailandensis* in DMSO-or RO-3306-treated cells. A549 cells were infected with *B. thailandensis* at MOI 100 and images were taken with a 20x objective between 15 to 17 hpi. Unpaired t-test with Welch’s corrections were performed (n=3, ns: not significant, * P<0.05).

**Figure S5.**
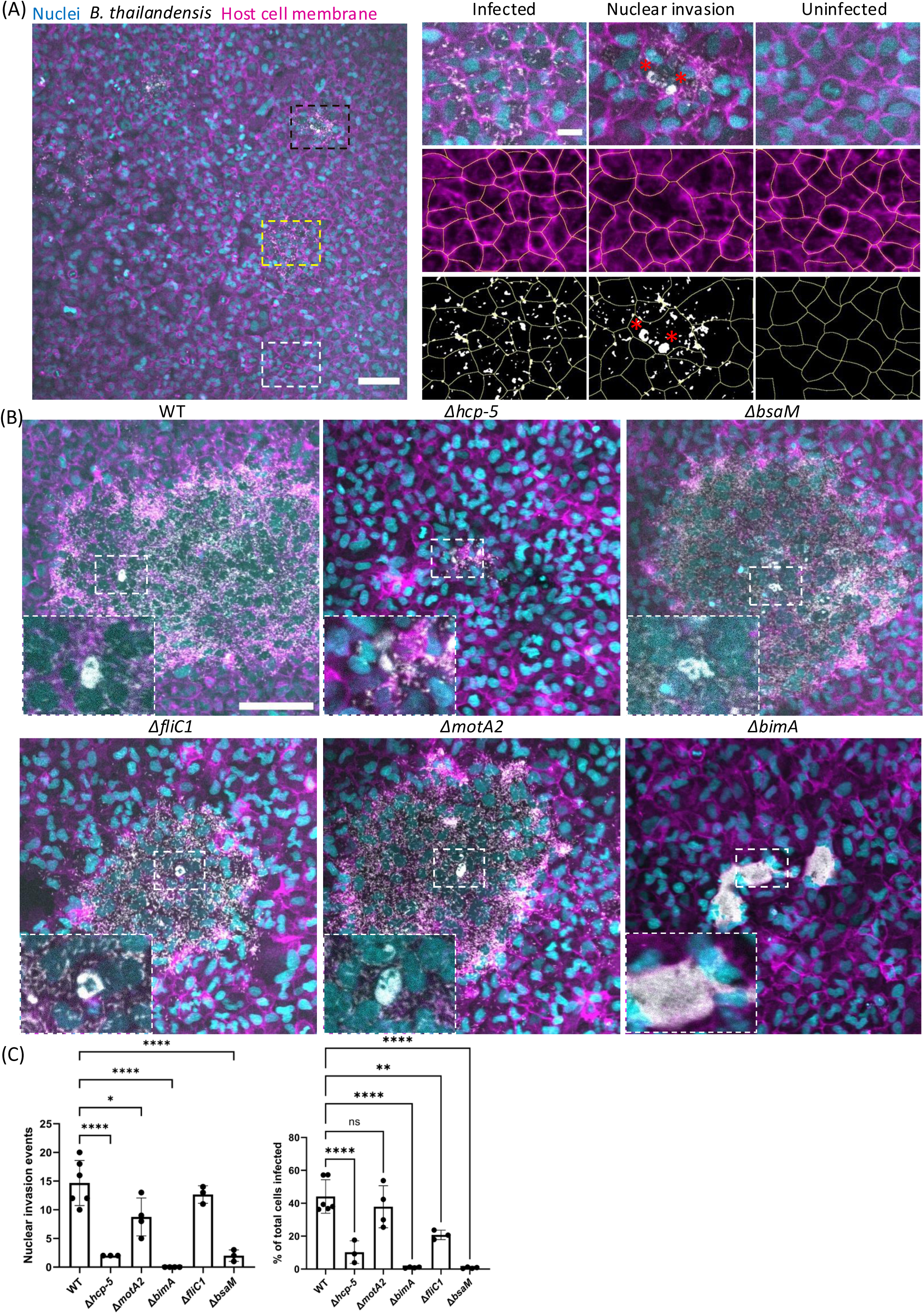
Examples of infected cell segmentation and nuclear invasion events. **(A)** A549 lung cells were infected with *B. thailandensis* expressing TssB-5-mScarlet-I (grey) at MOI of 100. Image was taken between 12:30 hpi with a 20x objective (905.54 μm x 905.54 μm). Host cell membrane was stained with CellMask DeepRed (magenta) and nuclei stained with Hoechst 33342 (cyan). Regions highlighted by yellow, black and white dotted lines represent the infected, infected cells with nuclear invasion, and uninfected cells respectively. In the magnified images, the top panel shows the raw image, middle panel displays eukaryotic cell segmentation with Cellpose, and bottom panel displays overlayed eukaryotic cell segmentation with Cellpose and fluorescent bacteria segmented with the Otsu algorithm. Scale bars represent 100 µm in the overview image and 20 µm in the magnified images. **(B)** Examples of various *B. thailandensis* mutants expressing TssB-5-mScarlet-I (grey) invading A549 cell nuclei at MOI 100. Host cell membrane was stained with CellMask DeepRed (magenta) and nuclei stained with Hoechst 33342 (cyan). Images were taken with a 20x objective between 14 to 17 hpi. Scale bar represents 100 µm. **(C)** Number of nuclear invasion events and percentage of total number of cells infected by various *B. thailandensis* mutants. A549 cells infected with various mutants of *B. thailandensis* at MOI 100 and images were taken with a 20x objective between 14 to 17 hpi. One-way ANOVA with Tukey’s multiple-comparisons test was used to determine the statistical significance (ns: not significant, * p<0.05, ** p<0.01, *** p< 0.001, **** p< 0.0001).

## Video legends

**Video S1. *B. thailandensis* TssB-5-mScarlet-I invades A549 cell nuclei, related to Figure 1**. Two examples of time-lapse images of A549 cells infected with TssB-5-mScarlet-I *B. thailandensis* at an MOI of 50. Images were acquired every 10 min from 5:30–17:50 hpi. T6SS-5 TssB-5-mScarlet-I protein fusion is shown in grayscale. Host cell membrane was stained with CellMask DeepRed (magenta) and nuclei stained with Hoechst 33342 (cyan). The time-lapse video is shown with 10 frames per second. The scale bars represent 20 μm.

**Video S2. T6SS-5 assembles in intranuclear *B. thailandensis,* related to Figure 2**. Videos showing examples of A549 nuclei infected with TssB-5-mScarlet-I *B. thailandensis*. T6SS-5 TssB-5-mScarlet-I protein fusion is shown in grayscale. Images were taken at 30 seconds intervals at the specified hpi. Host cell membrane was stained with CellMask DeepRed (magenta) and nuclei stained with Hoechst 33342 (cyan). The time-lapse video is shown with 5 frames per second. The scale bars represent 10 μm.

**Video S3. T6SS-5 assembles in cytosolic *B. thailandensis*** Δ***bimA,* related to Figure S2.** A549 cells were infected with TssB-5-mScarlet-I *B. thailandensis* Δ*bimA* at an MOI of 50. T6SS-5 TssB-5-mScarlet-I protein fusion is shown in grayscale. Images were taken at 30 seconds intervals at either 6:30 hpi or 13:00 hpi. Host cell membrane was stained with CellMask DeepRed (magenta). The time-lapse video is shown with 5 frames per second. The scale bars represent 10 μm.

**Video S4. Cytochalasin D treatment inhibits nuclear invasion events, related to Figure 7**. Time-lapse images of A549 cells infected with *B. thailandensis* expressing TssB-5-mScarlet-I (grey) at MOI of 50, and treated with cytochalasin D (5 µg/mL) or 0.05% of DMSO at 4 hpi. Images were taken at 15 minute intervals. Host cell membrane was stained with CellMask DeepRed (magenta) and nuclei stained with Hoechst 33342 (cyan). The time-lapse video is shown with 5 frames per second. The scale bars represent 10 μm.

